# TetrODrive: An open-source microdrive for combined electrophysiology and optophysiology

**DOI:** 10.1101/2020.12.16.423057

**Authors:** Marcel Brosch, Alisa Vlasenko, Frank W. Ohl, Michael T. Lippert

**Affiliations:** Systems Physiology of Learning, Leibniz Institute for Neurobiology, Magdeburg, Germany; Center for Behavioral Brain Sciences (CBBS), Magdeburg, Germany; Institute of Biology, Otto-von-Guericke University, Magdeburg, Germany

**Keywords:** microdrive, 3D printing, mouse, tetrode, fiber photometry, ventral tegmental area, optical tagging

## Abstract

**Objective:** In tetrode recordings, the cell types of the recorded units are difficult to determine based on electrophysiological characteristics alone. Optotagging, the use of optogenetic stimulation at the tip of electrodes to elicit spikes from genetically identified cells, is a method to overcome this challenge. However, recording from many different cells requires advancing electrodes and light sources slowly through the brain with a microdrive. Existing designs suffer from a number of drawbacks, such as limited stability and precision, high cost, complex assembly, or excessive size and weight.

**Approach:** We designed TetrODrive as a microdrive that can be 3D printed on an inexpensive desktop resin printer and has minimal parts, assembly time, and cost. The microdrive can be assembled in 15 minutes and the price for all materials, including the 3D printer, is lower than a single commercial microdrive. To maximize recording stability, we mechanically decoupled the drive mechanism from the electrical and optical connectors.

**Main results:** The developed microdrive is small and light enough to be carried effortlessly by a mouse. It provides high signal-to-noise ratio recordings from optotagged units, even across recording sessions. Owing to its moveable optical fiber, our microdrive can also be used for fiber photometry. We evaluated our microdrive by recording single units and calcium signals in the ventral tegmental area (VTA) of mice and confirmed cell identity via optotagging. Thereby we found units not following the classical reward prediction error model.

**Significance:** TetrODrive is a tiny, lightweight, and affordable microdrive for optophysiology in mice. Its open design, price, and built-in characteristics can significantly expand the use of microdrives in mice.

## Introduction

Extracellular unit recordings are an important tool in recording neuronal signals at single-cell resolution with high temporal precision. The spiking, recorded via metal electrodes inserted into the brain, has long been a key metric in advancing our understanding of the function of many brain areas, such as the hippocampus (MacDonald *et al* 2011), sensory cortex (Bao *et al* 2001), deep brain nuclei (Eshel *et al* 2015), and many more. Despite the existence of high-density silicon probes and recent advances in their long-term recording capabilities (Okun *et al* 2016), tetrode recordings are still irreplaceable due to their simple construction and high recording quality (Karumbaiah *et al* 2013, Harris *et al* 2000, Gray *et al* 1995). Multi-tetrode recordings combined with optic fibers in deep brain structures like the ventral tegmental area (VTA), locus coeruleus, or the amygdala are widely applied (Mohebi *et al* 2019, Eshel *et al* 2016, Takeuchi *et al* 2016, Zhu *et al* 2019). Traditionally, electrodes were implanted statically or mounted in stereotactic frames in acute experiments, limiting unit yield and awake recordings (Polikov *et al* 2005, Griffith and Humphrey 2006, Xie *et al* 2016, Nicolelis *et al* 2003). In contrast, head-mounted microelectrode drives (microdrives) allow recording in awake, head-fixed, or freely moving animals and repositioning of the electrodes (Chang *et al* 2013, Polo-Castillo *et al* 2019, Voigts *et al* 2020, Jackson and Fetz 2007, Santos *et al* 2012, Battaglia *et al* 2009).

Systems for use in rats typically allow the manipulation of dozens of independent electrodes (Ma *et al* 2019, Lu *et al* 2018, Billard *et al* 2018, Michon *et al* 2016, Kloosterman *et al* 2009). In mice, however, complex microdrives are challenging to use (Voigts *et al* 2013, 2020, Freedman *et al* 2016, Liang *et al* 2017, Rangel Guerrero *et al* 2018). Mice have a significantly smaller weight carrying capacity and allow only restricted microdrive dimensions, where even a few grams of weight already severely restrict the animal’s mobility. This effect becomes even more pronounced in juvenile mice (Battaglia *et al* 2009). Further, costs are high, and the assembly time of such intricate microdrives is long. For TetrODrive, we use desktop 3D printing to achieve a lightweight, stable microdrive that is assembled in minutes.

Integrating optical fibers into microdrives offers two main benefits (Liang *et al* 2017, Freedman *et al* 2016, Siegle *et al* 2011, Osanai *et al* 2019). Firstly, optogenetic stimulation can be used to alter behaviors or manipulate specific circuits (Kim *et al* 2017). Secondly, the stimulation can be used for optotagging, which confirms the cell identity of the recorded units by optogenetically eliciting spikes (Cohen *et al* 2012, Lima *et al* 2009, Mohebi *et al* 2019, Chen *et al* 2017). This allows a much more refined classification of the recorded cells; based on genetic cell identity rather than just waveform and discharge characteristics. If low-autofluorescence fibers are used, the fiber can be used to record fluorescence signals from the tissue around its tip via fiber photometry (Adelsberger *et al* 2005, Kim *et al* 2016, Cui *et al* 2014). Here we demonstrate fiber photometry recordings through the fiber built into TetrODrive.

Some microdrive designs for mice with an integrated fiber are built in a complex design with time-consuming assembly processes (Voigts *et al* 2013, 2020). Simpler single-screw driven designs for mice with optical fibers have been used (Anikeeva *et al* 2011, Wang *et al* 2018). However, the technical advancement of 3D printing allows rapid production and customization of microdrives. Previous 3D printed single-screw driven designs have a large footprint and immovable fibers (Kim *et al* 2020) or an assembly process with many different parts and increased weight (Delcasso *et al* 2018).

Building upon the principle of single-screw microdrives, TetrODrive is a novel, 3D printable, and easily assembled microdrive for mice, which combines up to eight tetrodes and one movable optical fiber in a lightweight design. The design is open and has been optimized to be 3D-printed with an affordable resin desktop 3D printer. The 3D printer itself costs less than a single commercial microdrive. In addition, our system addresses another issue of existing microdrives: mechanical coupling of plugging forces onto the electrodes. By separating optical and electrical connectors from the microdrive body, plugging and unplugging have no influence on the movable electrodes and fibers. Further, we demonstrate the use of our microdrive during long term recording of identified single units, optotagging as well as fiber photometric recordings in the VTA.

## Material and Methods

A list of all parts is available at the TetrODrive GitHub repository https://github.com/MarcelMB/TetrODrive.

### Tetrodes

Approximately 40 cm of tungsten wire (12.5 μm, HFV insulation, California Fine Wire Company) were twisted into a tetrode using an Arduino based twister (Open Ephys). The twisted wires were heat fused by a heat gun (temperature: 220 °C) from three sides. Eight of these tetrodes were attached to a custom-made electrode interface board (EIB, design files in the repository) with gold pins (Neuralynx). A thin layer of silicone was used as a conformal coating (3140 RTV Coating, Dow Corning Inc.). Following assembly, the electrode tips of the assembled microdrive were lowered into a cyanide-free gold plating solution (31 g/l, SIFICO ASC) and plated to 150 kOhm (plating current: −0.015 μA, 1 s, 30 runs, NanoZ, Multi Channel Systems).

### Optical fibers

Two different types of optical fibers were used: regular silica fiber for optogenetic stimulation (100 μm core, 0.22 NA, UM22-100, Thorlabs) and low-autofluorescence fiber for fiber photometry (200 μm core, 0.5 NA, FP200URT, Thorlabs). The fiber was connected to ceramic (optotagging) or steel (fiber photometry) ferrules (1.25 mm diameter) using UV glue (Loctite 4305). The fiber was inserted into a thin furcation tubing with Kevlar fibers (FT900KY, Thorlabs).

### 3D printing

The TetrODrive contains two 3D printed polymer parts: the main body (lower part) and the head (upper part). The TetrODrive was printed on a low-cost desktop printer using LCD technology (Photon, Anycubic). This printer hardens successive layers of resin (Cherry, HarzLabs) by transilluminating an LCD panel (50 μm resolution) with UV light (405 nm). The layer thickness was 50 μm. The layer cure time was 16 s for normal layers and 90 s for the six bottom layers. The pieces were printed orthogonally to the build plate and on thin supports. During one run (approximately two hours), the printer can print dozens of microdrive parts in parallel. Given the design of the parts, they can be printed on virtually any current resin printer. Filament-based (fused deposition modeling) printers are unsuitable, however, due to their low resolution. If no printer is locally available, online 3D print services can be used, or a milling machine can be used. Following printing, parts were briefly washed in isopropanol (< 1 min). It is critical to limit dwell time in the isopropanol as it facilitates stress cracking later on. Residual liquid was removed from the parts and holes using compressed air. The parts were post cured for 15 min under a UVA lamp and two hours of heating in an oven (50 °C). Design files, 3D files, and complete sliced printer files are available in the GitHub repository.

### Animals

All animal experiments were conducted according to the guidelines of the European Community (EUVD 6 86/609/EEC) and approved by a local ethics commission of the State of Sachsen-Anhalt. For electrophysiology and optotagging in combination with classical Pavlovian conditioning, we used four adult male and female (15–25 g) hemizygous C57/B6.SJL-Slc6a3^tm1.1(cre)Bkmn/J^ x C57/B6.129S-Gt(ROSA)26Sor^tm32(CAG-COP4*H134R/EYFP)Hze/J^ mice (Jackson laboratory, 006660 Dat-Cre crossed with 012569 Ai32). In these animals, Channelrhodopsin-2 (ChR2) is expressed under the control of the dopamine-transporter, labeling dopaminergic neurons (Tritsch *et al* 2014, Coddington and Dudman 2018). For fiber photometry, we used adult Dat-Cre mice (C57/B6.SJL-Slc6a3^tm1.1(cre)Bkmn/J^). All animals were housed under a twelve-hour light/dark cycle (light on at 6:00 AM). Food and water were provided *ad libitum,* except during the training period, when water was restricted to 1 ml per day Guo et al 2014), keeping body weight at around 85% of its initial value.

### Microdrive implantation

Anesthesia was induced by 3% isoflurane vaporized in air, followed by an intraperitoneal injection of sodium pentobarbital (60 mg/kg). Local anesthesia at the surgery site was provided by an injection of bupivacaine hydrochloride (0.25%, 0.2 ml). Mice underwent stereotactic surgery with implantation of a TetrODrive into the left VTA (AP: 3.5 mm, ML: −0.5 mm, DV: 3.9 mm). The microdrive was slowly (100 μm in one minute) lowered within a small craniotomy window (<500 μm diameter) until it reached its final position. Then, a silicone protection layer was applied around the insertion site, and dental cement was used to form a head cap. Two skull screws were implanted for stability and a custom-made head plate embedded in the dental acrylic. Two reference electrodes (chlorinated silver wires) were placed adjacent to the microdrive insertion site, and two silver ground wires were implanted above the occipital lobe. The EIB was fixated next to the microdrive, mechanically decoupling it from the microdrive body and allowing the plugging of the electrical and optical connectors without bending the drive mechanism and potentially disturbing the electrodes or fiber. Dental cement was applied until it covered most of the lower body of the microdrive. Care was taken to avoid dental cement reaching the metal cannulas or screw, as this would prohibit proper electrode movement. Parafilm^®^ was wrapped around the microdrive, tetrode wires, and EIB for protection. We found this to be a suitable lightweight alternative to a 3D printed enclosure in mice. Larger rodents may require a sturdier enclosure for the head-mounted parts. Animals recovered for at least one week.

### Fiber photometry

The mice underwent surgical procedures, as described above. Before inserting the microdrive, we injected 600 nl of an adeno-associated virus, carrying the fluorescent calcium indicator GCaMP6f (AAV.Syn.Flex.GCaMP6f.WPRE.SV40, 1E13 particles/ml, Addgene) into the left VTA (AP: −3.5 mm, ML: −0.5 mm, DV: 4.2 mm). Fiber photometry recordings started after three weeks of indicator expression. For recordings, the mouse was placed in the conditioning setup, and a custom-made, pre-bleached, low-autofluorescence fiber optic cable was connected to the ferrule. Excitation light for GCaMP6 was generated by a FiberOptoMeter (NPI Electronic), which also measured the returning fluorescence. The excitation light was modulated with a 66 Hz sine wave using a waveform generator (DG4102, Rigol), and the fluorescence signal was demodulated with a lockin amplifier (SR510, Stanford Research Systems) to reject ambient light changes. The resulting GCaMP6 signal was digitized at 1 kSp/s using a biosignal recording system (RHD2000, Intan Technologies) and analyzed in MATLAB (The MathWorks). The signal was detrended and filtered with a phase-neutral 20 ms sliding average filter to remove high-frequency noise. The fluorescence was calculated as ΔF/F, where F denotes the average pre-trial baseline fluorescence (1 s before trial start) and ΔF the difference between this resting light intensity and the ongoing signal.

### Electrophysiological recordings and spike sorting

Broadband signals (0.1 −20 kHz, at 30 kS s^-1^) of head-fixed mice were recorded via the RHD recording system (RHD2132 headstage, Intan Technologies) in a sound-attenuated and electrically shielded chamber. Spikes were detected and sorted using template matching with Kilosort2 (Pachitariu *et al* 2016a, 2016b). We used the default Kilosort2 configuration parameters, except for high-pass filtering, which was changed to 600 Hz, and spike-detection thresholds, which were reduced by half [5 2]. Bad channels were manually removed. Sorted spikes were manually curated using a python-based GUI for electrophysiological data (Cyrille Rossant, International Brain Laboratory).

### Electrophysiological data analysis

Data was analyzed with MATLAB (The MathWorks). We quantified the quality of the chosen single units using isolation distance (separation from neighboring clusters, measure for falsepositive spikes) and L-ratio (cluster spread, measure for false-negative events, Schmitzer-Torbert *et al* 2005). L-ratio threshold was 0.1 and isolation distance 20 for a cluster to be considered well isolated. Peristimulus time histograms (PSTH) were calculated using 1 ms bins. Histograms were convolved with function resembling a postsynaptic potential 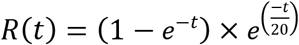 (Thompson *et al* 1996). Average spike waveforms displayed and used for cross-correlation calculation were band-pass filtered (300-6000 Hz). We calculated the peak width of the average waveforms of the highest negative amplitude deflection at half maximum amplitude. These points are more easily identified than the total spike length but naturally result in smaller absolute numbers. Non-identified units were only used for comparison in the session, where we also found optotagged units to ensure comparison in the same subregion. Each trial consisted of a conditioned stimulus (CS; tone) and an unconditioned stimulus (US; reward). CS-US ratios were calculated as 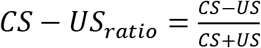. The CS and US ratio values are a difference between the peak firing rate (PSTH till 1 s after CS or US onset) and baseline firing rate.

### Optotagging

ChR2 expressing dopamine neurons were optogenetically identified by delivering short light pulses and analyzing the latency-locked, light-evoked spikes (Cohen *et al* 2012, Lima *et al* 2009). Laser light was provided by a blue diode laser (LightHUB-4^®^, Omicron). Before and after a behavioral session, light pulses (40 repetitions, 5ms duration, 473 nm) between 5-40 mW were delivered to elicit single opto-evoked spikes. The light intensity was adjusted to keep light-induced artifacts below a threshold of 1 mV and to ensure that the opto-evoked waveform was not distorted in comparison to the spontaneous spikes. We used various criteria to identify dopaminergic neurons, similar as it was applied previously (Cohen *et al* 2012): more than 50 percent of light pulses must evoke a light-induced spike, which has occurred in a time window of less than 10 ms post light onset. The cross-correlation of the spontaneous and light-evoked spike waveforms had to exceed r=0.9 to be considered as stemming from the same unit. The energy of the light-evoked spikes is defined as the integral of the squared voltage over time *E* = ∫ *v*^2^ *dt*. If we did not find optotagged units in a recording session, we advanced the microdrive more ventral by approximately 60 μm and recorded on the following day.

### Behavioral task

Water-deprived mice were handled and habituated to the head-fixation setup for three days without experiencing any reward in the setup. During the head fixation, mice could run on a circular treadmill. Following habituation, the mice underwent a Pavlovian, auditory trace-conditioning task. The trial structure consisted of a CS (pure tone, 60 dB, 5 ms onset/offset ramp, 500 ms, 16 kHz) delivered free-field (UltraSoundGate Player 116, Ultrasonic Dynamic Speaker Vifa, Avisoft Bioacoustics) and a 1.5 s delayed US (water reward, 5 μl, 2 s delay for fiber photometry). The tone was quieter than salience tones that elicited dopamine neuron salience responses in primates (Fiorillo *et al* 2013). The water was dispensed by a solenoid valve (The Lee Company), placed outside the sound attenuated setup in its own sound absorbing enclosure. The lick spout was a blunted syringe needle. Licks were detected through the breakage of an infrared beam in front of the lick spout. If the animal licked less than one second before tone onset, the trial was aborted, and a new trial was initiated. This procedure prevents random licking. The intertrial intervals were computed from a randomly permuted exponential distribution (mean of 10 s, for fiber photometry 20 s). One session consisted typically of 100 to 150 total trials. In ten percent of trials maximum, the US was omitted, and in an additional maximum of ten percent, the US was given without a CS. In selected trial blocks with incorporated CS and US omitted trial types, these trial types never exceeded 30% and never occurred consecutively. If animals received less than 1 ml water within the session, additional water was given after the session in the home cage to maintain the desired body weight. We controlled our behavioral setup with an Arduino, running custom code based on previous work (Micallef *et al* 2017). Responses were visualized live using a custom MATLAB (The MathWorks) interface. Timestamps generated by the Arduino were recorded by the RHD2000 recording system to synchronize behavior and electrophysiology.

### Histology

At the end of the experiment, animals were killed with an overdose of pentobarbital and perfused with formaldehyde in 0.1 M phosphate-buffered saline (4%). Brains were extracted and cryosectioned into 50 μm coronal slices. Slices were mounted in polyvinyl alcohol mounting solution (Mowiol ®), and fluorescence images of the recording sites in the VTA were taken.

## Results

### Assembly of TetrODrive

TetrODrive consists of only two 3D printed parts, and its assembly takes approximately 15 minutes (figure 1(a)). This process requires the following steps: the middle hole of the printed microdrive body is widened with a 0.8 mm drill, followed by tapping for M1×0.25 thread (figure 1(b)). Tapping must be done carefully to avoid destroying the fragile threads. The drill should be repeatedly removed to clear forming resin debris from the tap. The two holes holding the cannulas in the body must be widened completely with a 1 mm drill to achieve a precise gliding motion (figure 1(b)). It is beneficial to test the drilling result with a piece of later used cannula for smooth movements. All drilling should be done by hand, holding the drill in one hand, clamped into a small manual vice, and the microdrive part in the other.

**Figure 1.**
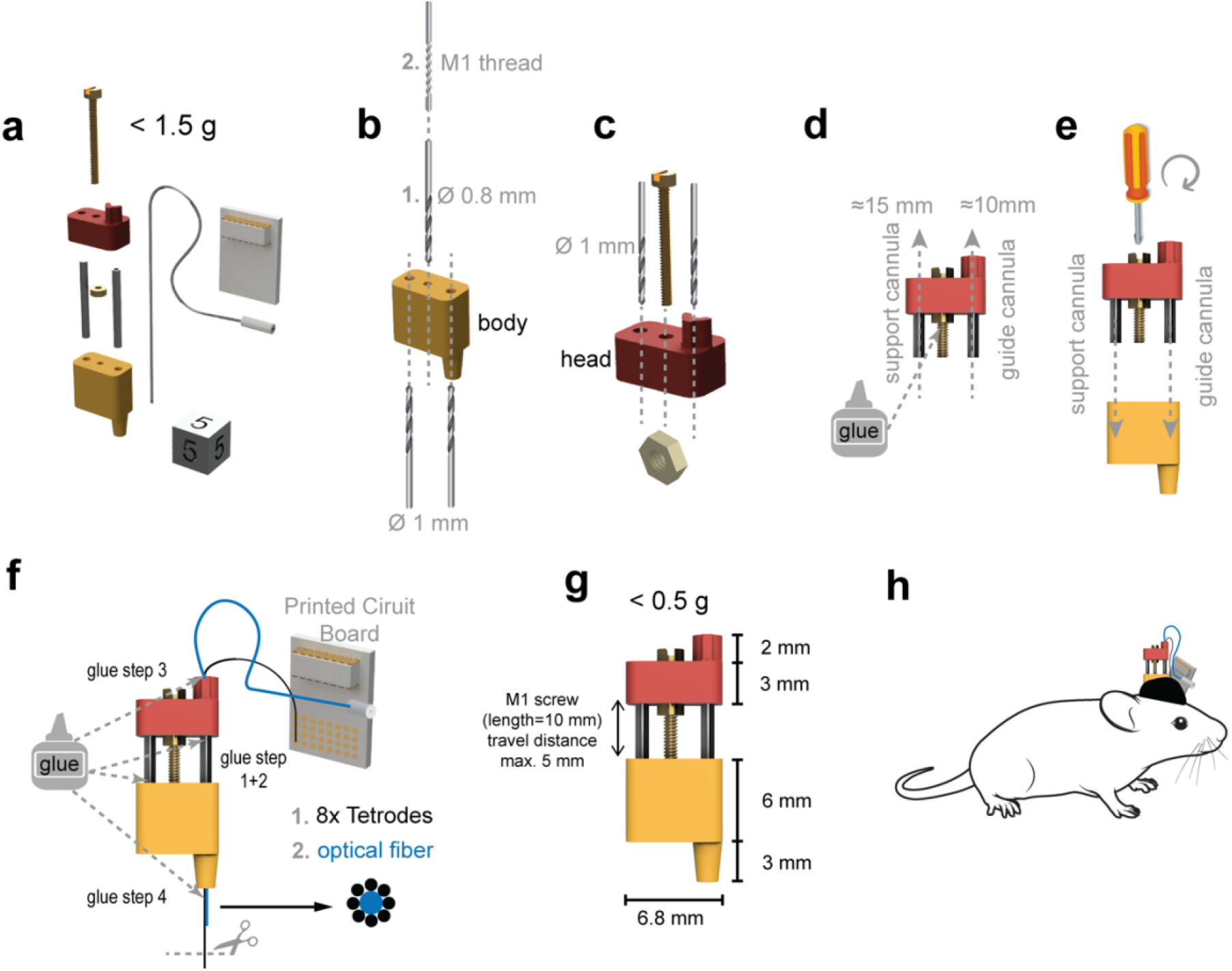
Assembly and loading of the TetrODrive. (a) The TetrODrive consists of only two 3D printed parts, one screw, a nut, and two steel cannulas. Simple construction allows it to be assembled and loaded in less than 15 min. (b) Holes in the body part are drilled to fit a 1 mm cannula size, and a thread is cut for the screw. (c) Holes in the microdrive head are also enlarged. A screw is inserted and fixed with a nut from the bottom. (d) The nut is fixed with glue to the screw and cannulas are inserted from the bottom. (e) The microdrive head is connected to the body and the screw is turned a few turns. (f) The guide cannula for fibers and tetrodes is glued to the head and the support cannula was glued to the body part. Tetrodes and fiber are glued to the protrusion on the microdrive head. The tetrodes are fixed with glue to the fiber. Tetrodes are cut to length with sharp, carbide-tipped scissors and later gold plated. (g) Dimensions of the TetrODrive. The length of the screw determines the dorsoventral travel distance. (h) Illustration of the TetrODrive implanted in a mouse. The microdrive complete body part and the EIB lower end are embedded in dental cement but mechanically isolated from each other.

Following the microdrives body’s preparation, the microdrive head is assembled: First, the two outer holes are widened to 1 mm. Then, a 10-millimeter long M1 screw is fastened into the microdrives head’s middle hole and a nut is fastened against the lower side of the microdrive head (figure 1(c)). The nut must be tight enough to prevent wiggling, but it must not unduly restrict the turning of the screw. The nut is then fixed to the screw with a tiny drop of UV glue (figure 1(d), Loctite 4305 UV glue). Stainless-steel cannula tubing (1 mm diameter, hollow inside) is prepared for the 10 mm long guide cannula and 15 mm long support cannula. All tetrodes and the fiber sit inside the guide cannula. Cannula ends must be carefully deburred to prevent tetrode damage. Both cannulas are inserted into the microdrive head (figure 1(d)). We chose affordable and readily available materials for our implementation. Alternative materials can also be used to accommodate non-metric materials. The length difference between the guide and support cannulas is approximately the drivable distance for the TetrODrive (in this case, five mm).

The microdrive head with its guide and support cannulas is inserted into the microdrive body, and the screw is carefully turned a few turns into the microdrive body, joining the two subassemblies (figure 1(e)). Using a small amount of UV glue, the support cannula now has to be secured in the microdrive body and the guide cannula in the microdrive head (figure 1(f), glue step 1+2). This allows the guide cannula fixed to the head to move up and down using the fixed body support cannula. By turning the screw, thereby moving the head up and down, the correct assembly is confirmed.

The bottoms of the support cannula hole and the tapped screw hole should be blocked with a piece of tape against the ingress of fluids or glue during surgery. The dimensions and low weight of the microdrive are designed for mice, including juvenile mice used in this study (figures 1(g), (h)).#

### Tetrode and fiber loading

Tetrodes connected to an EIB are inserted into the guide cannula until they protrude a few millimeters beyond the planned initial implantation depth from the lower side of the microdrive body (figure 1(f)). Care should be taken not to damage the tetrode insulation. Following tetrode loading, the optical fiber is inserted into the guide cannula until it protrudes the planned initial implantation depth. The fiber and tetrodes are then secured to the guide cannula at the microdrive head using UV glue (figure 1(f), glue step 3). The protruding electrodes are UV glued to the fiber and cut to length using carbide-tipped scissors (regular steel scissors are too soft and can lead to poor tetrode performance, shorts and coating crosslinks, figure 1(f), glue step 4). We kept the distance between fiber and electrode tips in the range of 300 to 500 μm. Ideally, tetrodes should be equally spaced around the fiber tip and not clustered in one spot. UV glue must not touch the fiber’s tip, and only small amounts should be used to keep the implant diameter low. In the last step, the ferrule at the end of the fiber is glued to the backside of the EIB (figure 1(f)). The assembled microdrive is now ready for tetrode plating, sterilization, and implantation. Design files of the TetrODrive and EIB are available in the repository.

### Chronic single-unit recordings

We recorded single units from tetrodes implanted in the VTA. Exemplary units, sorted in Kilosort2, are depicted in figure 2(a). Auto-correlograms show a clear refractory period, and crosscorrelograms no common refractory period (figure 2(b)). An average of 5 ± 1 (mean ± SEM) well-isolated units per tetrode and recording session (n= 3 animals, 7 sessions) were found. Units could be recorded for many weeks, particularly if the tetrodes were continuously advanced (figure 2(c)). To test long term recording quality without electrode movement, we recorded from a preparation in which the microdrive had not been moved for six weeks and implanted for ten weeks. Even under these conditions, we still found clear single units, showing high signal-to-noise ratio spikes with amplitudes in the millivolt-range (figure 2 (d)). We find that, if units are lost, after moving tetrode tips forward for about 100 μm (1/2 turn), many new units will appear in the next recording (figure 2(e)). We found it beneficial to advance the electrodes a day before recording, allowing the tips to settle. Particularly due to the relatively thick glass fiber, tissue movement can require some time (Bakhurin *et al* 2020). However, this settling period may vary with implantation depth, fiber thickness, and structure of interest and should be experimentally determined for each preparation.

**Figure 2.**
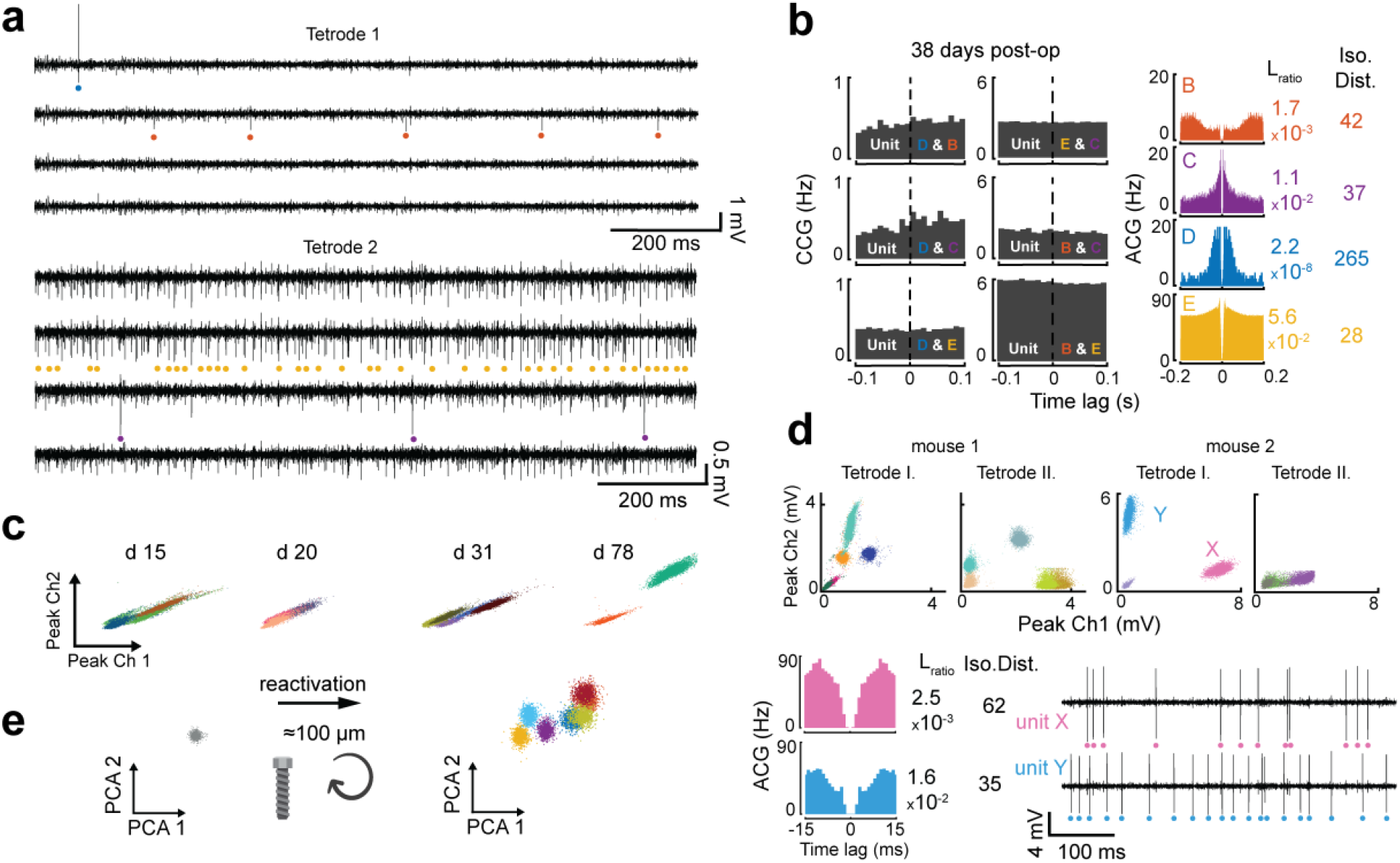
Chronic single-unit recordings with and without microdrive movement (a) Exemplary, a band-pass filtered signal from two tetrodes. Unit spikes are color-coded. (b) Unit quality. Only units with low L-ratio and sufficient isolation distance (iso. dist.) were included. Autocorrelations (ACG) for each unit and cross-correlations (CCG) between units (38 days after implantation) demonstrate unit separation. (c) Peak amplitudes of two channels from the same tetrode over time. Discrimination of units during four different time points with microdrive movement between recording sessions. (d) Exemplary remaining single units in the absence of microdrive movement after 42 days in one position (72 days post-op). (e) Tetrodes which do not show active units after a prolonged time (1 unit / 3 tetrodes) at one position can be reactivated by turning the microdrive screw (7 units / 3 tetrodes). (n=3)

### Single-unit stability and fine depth adjustments

The fine thread of the used screw and the mechanical decoupling allow precise movement of the electrode tips in the brain and resistance to plugging forces. Figure 3(a) depicts the identical two units from two tetrodes as figure 2(b) but shows their evolution across recording sessions and drive motion. 180° screw turns (> 125 μm) result in new sets of single units (figure 3(a), position 1, rec. 1 – position 2, rec. 2; position 3, rec. 6 – position 4, rec. 7). For five consecutive recordings sessions, these units could be followed (figure 3(a), rec. 2 – rec. 6, r> 0.9). For example, unit B shows high intra-session and inter-session amplitude stability (figure 3(c) rec. 3 – 4, position 2). A stable coefficient of variation for interspike intervals (C_v_) was an additional measure for unit stability over the recording session (e.g., unit B_1-5_ and E_1-5_, figure 3(a)). These results demonstrate that TetrODrive can be used to record stable units across multiple days, even if the fiber is plugged and unplugged several times.

**Figure 3.**
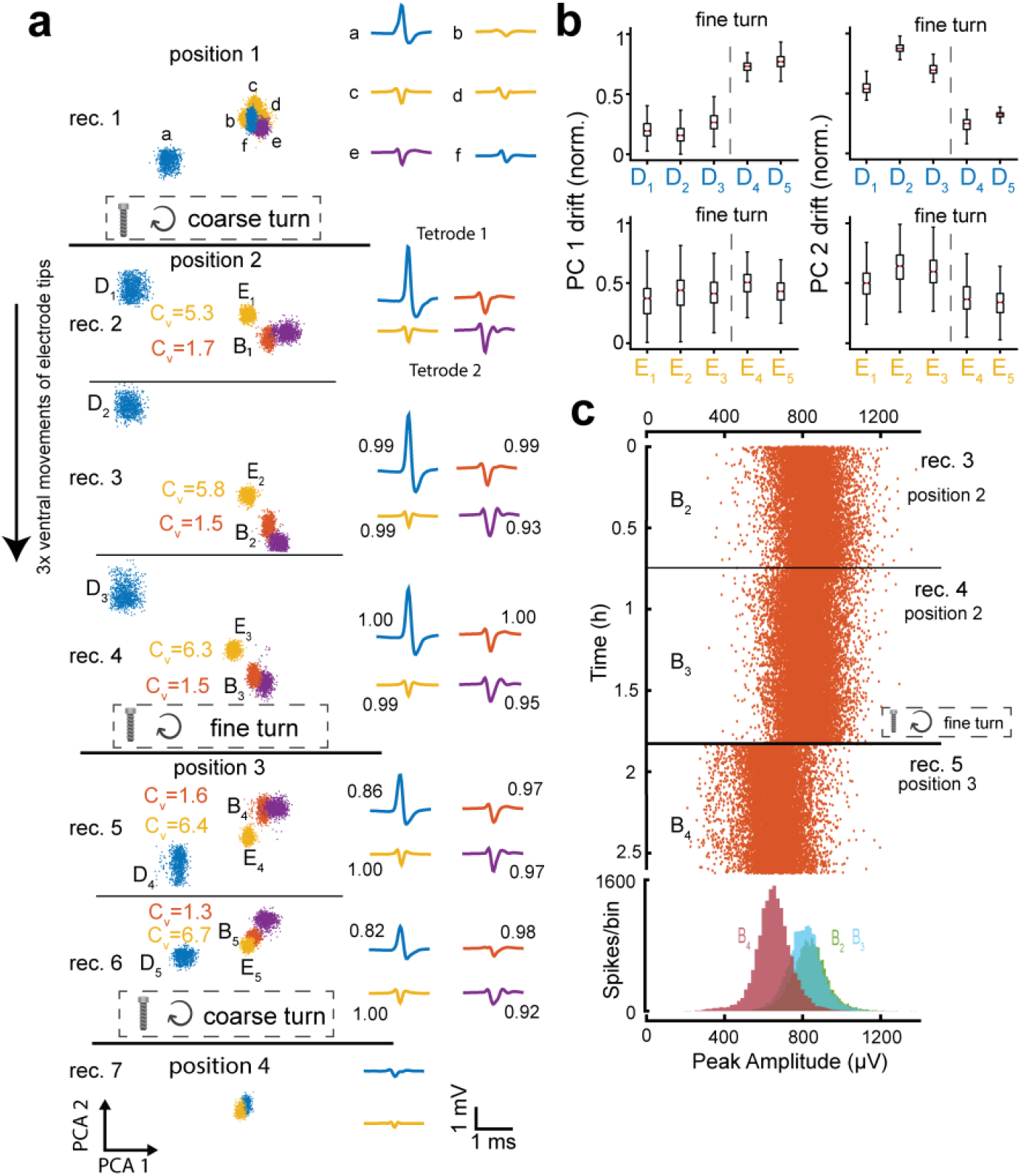
The precise driving mechanism of the TetrODrive. (a) Single unit stability over consecutive recording sessions (rec.) and unit changes due to microdrive movement on two exemplary units for two tetrodes. Principal components 1 and 2 (PC) and mean waveform (with cross-correlation of the previous recorded similar unit for 5 recording sessions) of exemplary units at four different positions. New units were acquired due to depth adjustments of more than 125 μm (coarse turn) from position 1 to 2. Units from position 2 were stable over three consecutive recording sessions. A fine adjustment of less than 60 μm (fine turn) resulted in a shift of the same units from position 2 to 3. New units were acquired again due to depth adjustments of more than 125 μm (coarse turn) in position 4. (b) Drift of PC 1 and 2 of unit D and unit E due to fine (90°) microdrive turn. (c) Spike amplitudes from unit B for three consecutive sessions (recording sessions 3 to 5). A fine adjustment resulted in a slight shift of the spike peak amplitudes on the best channel. (n=1)

Fine screw adjustment not more than 90° (< 60 μm, position 2, rec. 4 – position 3, rec. 5) move the electrodes so little, that most units can still be followed with only slight shifts in mean waveform (figures 3(a)). Waveform correlations remain above r>0.9 for most units for 5 days (rec. 2 – rec. 6). The fact that the tip did indeed moved is demonstrated by the change in unit D (r>0.8, shift in PC components, figure 3(b), unit D upper panel) and unit B amplitude change (figure 3(c), position 2, rec. 4 – position 3, rec. 5). For unit E, this shift in PC components is more subtle and only observed in the PC 2 (figure 3(b), lower panel). These results demonstrate that the TetrODrive is stable enough to keep units across days but can move precisely enough to allow fine, continuous movement through the tissue, which is helpful once optogenetically responsive neurons are found and need to be followed.

### Optogenetic identification of dopaminergic midbrain neurons

Here we demonstrate the ability of TetrODrive to perform chronic optotagging in the VTA during Pavlovian conditioning. We expressed ChR2 in dopaminergic cells (figure 4(e)) and optogenetically stimulated these cells with blue laser pulses. In dopaminergic cells, this stimulation leads to a single, short-latency spike after light-onset (figure 4(a)). Characteristically, such neurons can follow only low stimulation frequencies (figure 4(b)). We plotted the correlative relationship between average spontaneous waveform (figure 4(c), black waveform) and average optogenetically evoked waveform (figure 4(c), blue waveform) against the light-evoked voltage response, which revealed responsive (blue) and non-responsive (gray) units (figure 4(d)). Interestingly, we observed units which showed changes in their response latency to the laser pulse, when comparing it between before and after Pavlovian conditioning (unit 1, figure 4(c)). Identified dopaminergic units had an average firing rate of 4.8 Hz (range: 1.9 – 9.4 Hz). Mean peak width was 151.6 μs (range 81.2 – 261.30 μs, full width half maximum). Optogenetically non-responsive single units, in sessions which had at least one confirmed dopaminergic unit, had higher mean firing rates of 24.9 Hz (range: 0.5 – 71.7 Hz) but similar mean peak width of 150.5 μs (range: 81.8 – 401.45 μs).

**Figure 4.**
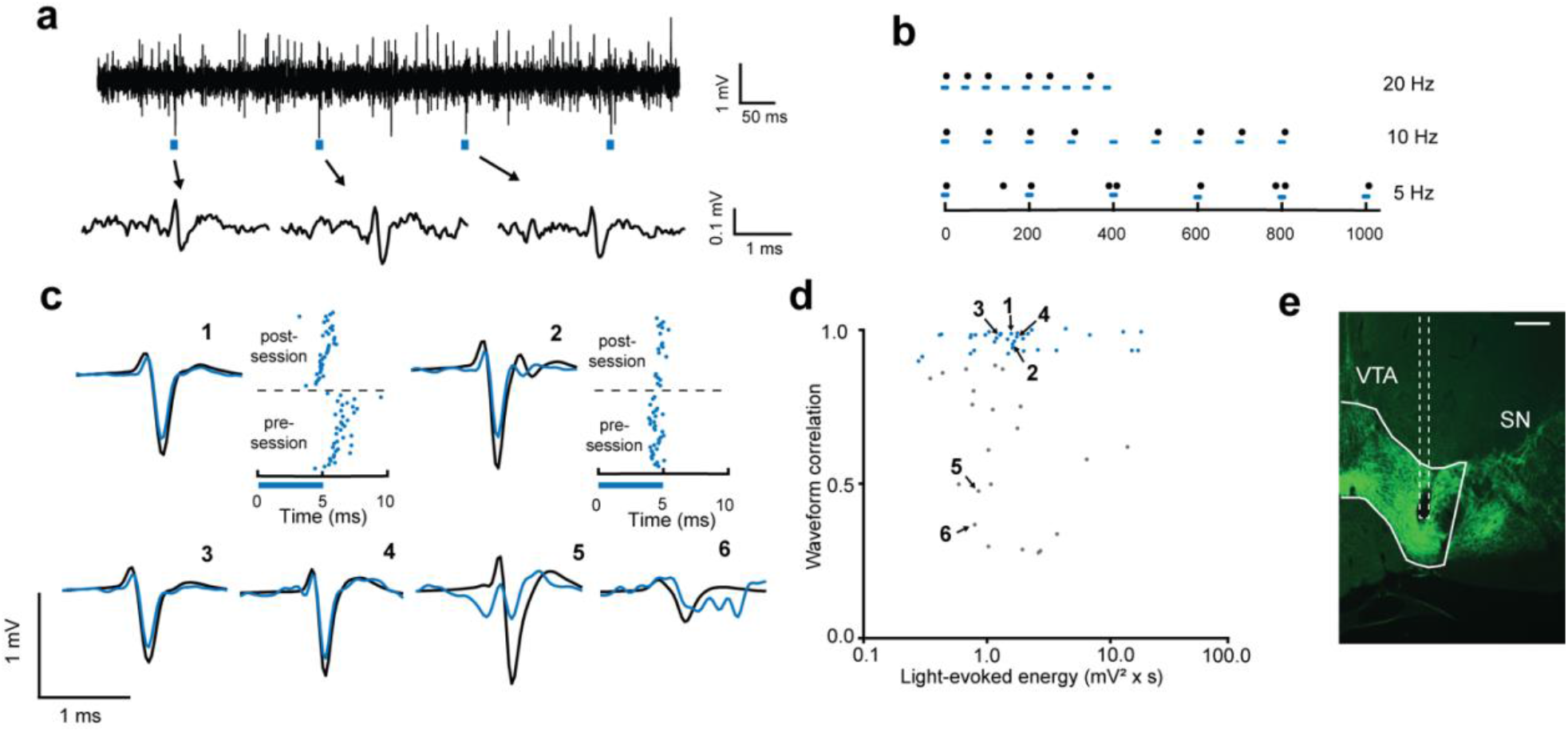
Identifying dopaminergic neurons in the VTA (a) Continuous band-pass filtered signal. Short light pulses (blue square, 5 ms) elicited a single time-locked light-evoked spike. (b) Response from an optotagged neuron for 5, 10, and 20 Hz light stimulation. Black dots represent a spike; blue squares represent a light pulse. (c) Correlation of spontaneous spike waveforms (black) and light-evoked waveforms (blue) before and after conditioning sessions. Raster plot of spikes around laser onset (0 ms) before and after conditioning. The onset latency of identified units has shifted during conditioning. (d) Cross-correlation of spontaneous and light-evoked waveform against the light-evoked spike energy. (e) Exemplary histological slice of fiber and tetrode position in the VTA (green: Dat specific ChR2-eYFP expression). Anatomical structures based on Paxinos and Franklin (2001). Scale bar=200μm, (n=4)

### Reward prediction error in midbrain dopaminergic neurons

Following previous work, we recorded the change in response to anticipated and surprising US in the VTA (Schultz *et al* 1993, Cohen *et al* 2012, Roesch *et al* 2007, Bayer and Glimcher 2005). Recording of the same dopaminergic single unit over four consecutive conditioning sessions showed an increased response strength to CS but a more diverse response strength to anticipated US (figure 5(a)). After the presentation of an unpredicted US, dopaminergic neurons responded strongly (figure 5(b)). The omission of a predicted US resulted in a firing rate decrease at the time of the anticipated US, best observed in the raster plot (figure 5(c)). We generally observed high diversity of identified dopaminergic unit responses, with some units responding strongly to the US, to the CS, or both, even after extended conditioning (figure 5(d), (f)). Despite the observed response-diversity of dopaminergic units, overall, we observed a shift of response strength from the US towards CS over conditioning sessions (figure 5(e), r = 0.66, P<0.001). The increase in CS response was less pronounced (r = 0.52, p<0.01) than the decrease in US response (r = 0.28, p=0.17, figure 5(f)).

**Figure 5.**
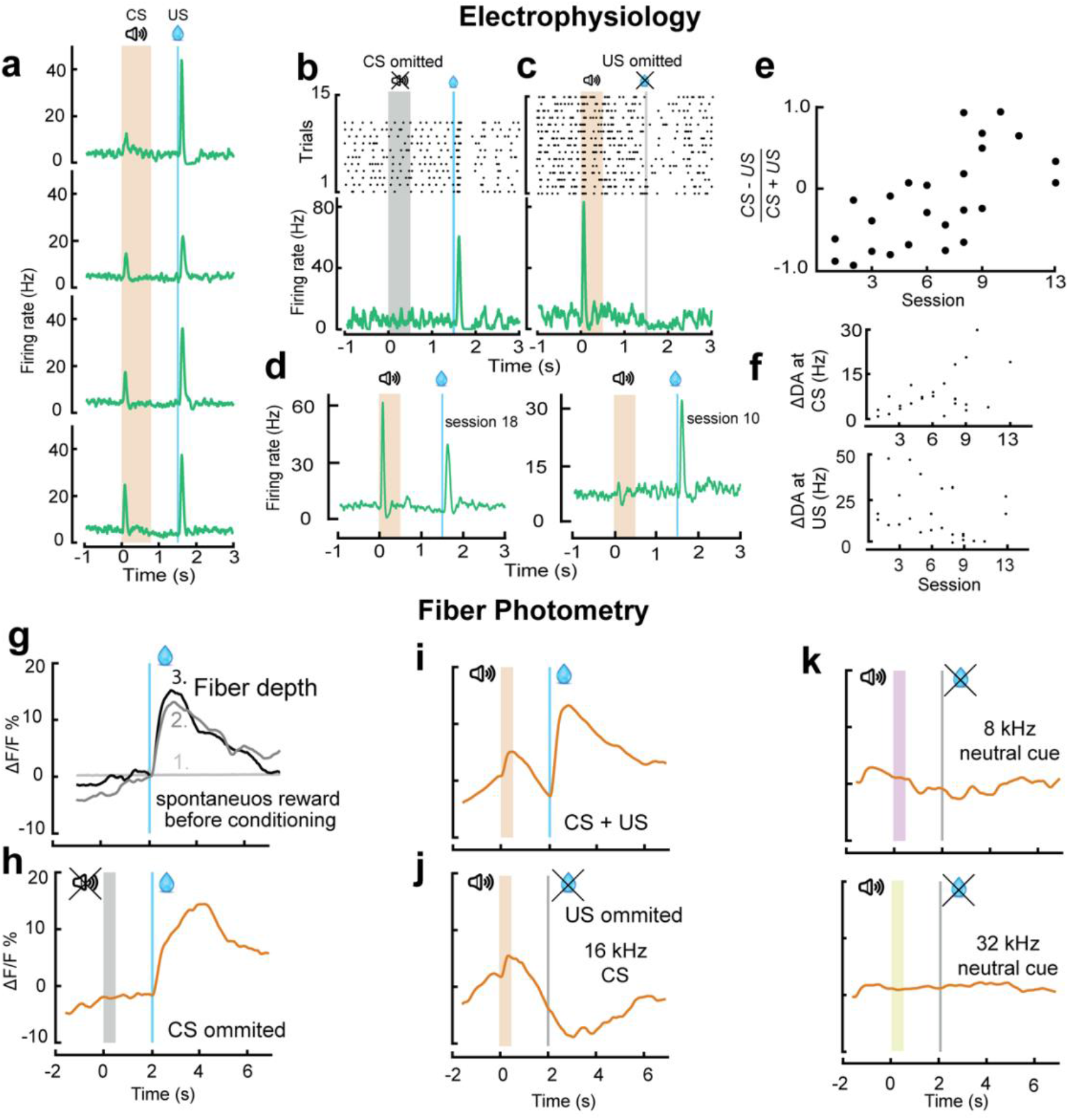
Dopamine reward prediction errors during Pavlovian conditioning. (a-f) Single unit recordings of optotagged dopamine neurons (n=3). (g-k) Genetically encoded dopaminergic bulk fluorescence through fiber photometry (n=1). (a) Average session responses of an exemplary dopaminergic unit over three consecutive sessions. The response to the CS increases over time, whereas the response to the US is not extinguished. (b) Positive reward prediction error response can be seen following the unpredicted US. (c) Following extensive training, cells respond to the CS tone but show a negative reward prediction error at the time of an omitted US. (d) Average session response of dopaminergic units in two late behavioral training sessions. (e) CS-US ratios for dopaminergic units over the behavioral training. (f) The average session response firing rate (minus baseline activity) for dopaminergic units at the time of the CS and US as a function of behavioral sessions. (g) Adjusting the fiber tip position can be used to optimize fiber photometry signal amplitude. (h) Positive reward prediction error in trials with unpredicted US in a trained animal. (i) Response to the CS and the US in a trained animal. (j) Negative reward prediction error in trials with a CS, but omitted the US in a trained animal. (k) Response to two neutral cues.

For fiber photometry, we replaced the regular optical fiber in our TetrODrive with a low-autofluorescence fiber. At the initial implantation depth, above the VTA, no reward-related fluorescence response could be detected. Response magnitude to unexpected water rewards increased the further ventral the fiber was moved (figure 5(g)). Once we reached an optimal depth for fluorescence recording, we started the Pavlovian reward conditioning task. In this task, we found a response to the US, both unexpected and expected, and to the CS (figure 5(h), (i)). For the omitted US, we observed a phasic decrease in fluorescence (figure 5(j)). These findings are compatible with reward prediction error coding. The response to the US was present even after many conditioning sessions. We did not find any response for neutral sounds not paired with a reward (figure 5(k)).

## Discussion

Here we present a novel, 3D printed microdrive for combined electrophysiology and optophysiology in mice. It can be manufactured from few parts at extremely low cost in virtually any laboratory, is highly compact, lightweight, and provides mechanical decoupling of all connectors from the movable microdrive parts. A movable fiber allows optogenetic stimulation as well as fiber photometry. All design resources are openly available.

Larger rodents allow the implantation of relatively large microdrives, and as a result, most existing designs target rats instead of mice (Lu *et al* 2018, Ma *et al* 2019, Keating and Gerstein 2002, Kloosterman *et al* 2009, Lansink *et al* 2007, Billard *et al* 2018). For mice, a smaller number of microdrives exist, but none so far have combined the features of the TetrODrive. Typically, optogenetics-enabled 3D printed mouse microdrives fall into two categories: A) large, complex and heavy microdrives that allow the individual manipulation of each electrode (Voigts *et al* 2020, Freedman *et al* 2016, Liang *et al* 2017, Brunetti *et al* 2014). They are advantageous for experimental approaches that require all electrodes to target a relatively thin cell layer, such as in the hippocampus; B) simple microdrives that are smaller, do not allow manipulation of individual electrodes and typically lack movable fibers (Kim *et al* 2020, Headley *et al* 2015). The complexity and weight of a simple microdrive designs increases by incorporating movable fibers (for example, up to 4.4 g), causing limitations to work with young mice (Delcasso *et al* 2018). Therefore, other simple designs include only a static fiber (Kim *et al* 2020). The TetrODrive is the first 3D printed simple design multi-tetrode microdrive with a movable optic fiber, that is single screw driven and lightweight (0.5 g).

Integrating optical fibers into microdrives can cause another problem: the optical connector is typically attached directly to the assembly’s driven part, coupling mechanical forces onto the electrode tips during plugging cycles. The problem has been recognized, and solutions were developed previously for microdrives without fibers and silicon-probe microdrives (Ma *et al* 2019, Chung *et al* 2017, Freedman *et al* 2016). The design of the TetrODrive, splitting drive mechanism and connectors, inherently avoids the coupling of plugging forces onto the electrodes and fiber. To further reduce forces, we use zero insertion force connectors, in which almost no net plugging force is exerted onto the head. As a result, we could record from the same optotagged units for multiple consecutive sessions, while previous studies advanced electrodes after each recording session (Eshel *et al* 2015).

Our microdrive does not feature individually moveable tetrodes. This may limit its usefulness for preparations in which several tetrodes need to be positioned in a thin cell layer, such as the hippocampus (Ulanovsky and Moss 2007, Pfeiffer and Foster 2013). We consciously made this choice to reduce the complexity of the microdrive and its assembly process. Microdrives with individually movable electrodes can easily take days to assemble while our design is assembled within 15 min. In brain areas that do not require relative movements between individual electrodes, it is unnecessary to invest this time and have the animal carry the excess weight and bulk.

To demonstrate the versatility of TetrODrive, we employed it for recording, optotagging, and fiber photometry in the VTA. The VTA is a brain area strongly implicated in the processing of reward but reveals a functionally diverse dopaminergic activity pattern (Bromberg-Martin *et al* 2010, Roeper 2013, Schultz *et al* 2017). It is composed of different neuronal subpopulations (Lacey *et al* 1989, Nair-Roberts *et al* 2008), making optotagging instead of electrophysiological markers a necessity to reliably identify the recorded units (Ungless and Grace 2012, Margolis *et al* 2006).

In general, our data is in alignment with the reward prediction error theory. The responses to the anticipated US gradually weaken, and responses to the CS grow. We observed, however, inhomogeneity in responses from the identified dopaminergic units. Many units still strongly responded to the US, even after conditioning, in accordance with previous observations (Cohen *et al* 2012, Coddington and Dudman 2018). This finding could be explained by sensory or motor processes influencing the firing (Coddington and Dudman 2019, Howe and Dombeck 2016, Da Silva *et al* 2018, Hughes *et al* 2020). A second explanation for sustained responses to the US could be reward prediction error coding with long eligibility traces (Pan *et al* 2005).

In contrast to previous microdrives, our design also allows fiber photometry and the optimization of the recording location through the moveable fiber. In the population calcium signal that we recorded from the dopaminergic cells of the VTA, we observed some hallmarks of reward prediction error coding, such as responding to the CS and attenuated responses to omitted US. Again, we did not find a cessation of responding to anticipated US, even after prolonged conditioning. However, the combination of fiber photometry and electrophysiology demonstrates for the first time that this property is characteristic for the VTA on a population level but not present in all individual units. Therefore, results from fiber photometry should therefore not be taken as representative for all units in a particular area but interpreted with caution.

Future iterations of TetrODrive could be improved by integrating a remote drive mechanism. Using a miniature motor (Yamamoto and Wilson 2008, Fee and Leonardo 2001) or piezo crystals (Smith *et al* 2020), the microdrive could be shielded from all external mechanical forces. This might help to increase unit stability even further. Other avenues for development, which we have partially tested already, would be to further miniaturize the microdrive. Due to its simple construction and use of off-the-shelf parts, smaller screws and cannulas can be used to save even more weight and volume. Currently, approximately two microdrives can be used per mouse, especially if the EIB is shared between microdrives. With a smaller design, more microdrives could fit (Headley *et al* 2015).

## Conclusion

TetrODrive is a 3D printable, easy to assemble, and open microdrive design that combines optophysiology and electrophysiology. Its unique combination of simple design and advanced performance opens the use of microdrives for mice to many more laboratories. By combining electrophysiology and optophysiology, it also bridges the gap between traditional unit recording and genetic methods, such as optogenetics and fiber photometry.

## Acknowledgments

This work has been supported by the Center for Behavior Brain Science Magdeburg (CBBS, FKZ ZS/2016/04/78120), a German Research Foundation Grant (Priority Program 1665 of the DFG/OH 69/1-2) and the Leibniz Society (LIN Postdoc Network). The authors declare that the research was conducted without any commercial or financial relationships and therefore state no conflict of interest.

We thank the Uchida lab at Harvard University for VTA recording and optotagging advice and inspiration in the conception of TetrODrive. Timothy Michael French helped us with manuscript corrections. We thank SciDraw.io for providing a mouse vector graphic that was adopted in figure 1(h) under the creative common license (CC BY 4.0).

